# Reaching movements in virtual reality depend on the seen avatar arm and not just the acting arm

**DOI:** 10.64898/2026.10.05.756878

**Authors:** Olivia Ipwanshek, Karsten Babin, Paul L Gribble, Jody C Culham

## Abstract

Virtual reality (VR) offers unique opportunities to study behaviour by allowing the visual environment to be manipulated in ways otherwise impossible in the real world. A central theoretical question is how body-related visual information shapes movement control. Given right-handers’ greater dexterity with the right arm, perhaps even seeing a right arm vs. a left arm may engage sensorimotor processes associated with more precise motor control. We examined reaching performance while participants used their own right or left arm to move the manipulandum of a KINARM robot to intercept static targets. Participants wore a VR head-mounted display, calibrated to the robot’s physical workspace, in which they saw either the right or left arm holding and moving the manipulandum. The 2 × 2 design allowed for independent manipulation of the controlling arm and virtual arm displayed. As expected, reaches were faster and more accurate when controlled by the right arm. More interestingly, performance was also affected by the seen arm. Participants made straighter reaches when they saw a virtual right arm than a left arm, especially for the right controlling arm. These findings show that precise motor control relies not only on the arm that is actually moved and provides proprioceptive feedback, but also on the visual identity of the arm. The ability to modulate performance using visual features in VR opens new theoretical questions and opportunities for applications such as rehabilitation.

## INTRODUCTION

We use vision to guide motor behaviour, such as reaching for a cup of coffee or writing with a pen. These actions depend on a continuous interplay between motor commands and sensory feedback. As we move, both vision and proprioception provide information about the position and movement of the body, with vision relying on visual input and proprioception on sensory signals from muscles and joints. Together, these sources of feedback allow movements to be adjusted during action (Chiel & Beer 1997; Wolpert & Flanagan 2001, 2010). Understanding how sensory information contributes to action is therefore critical for explaining how the brain plans, executes, and refines goal-directed movements.

Studies of the motor system often investigate visually guided reaching using simplified visual feedback, such as a cursor representing hand position (Krakauer 2019; Saunders & Knill 2003; Saunders & Knill 2004). Although cursor-based tasks are useful, they also remove many body-specific cues that are present during real-world action. In real-world action, we often see our own body moving through space. When we reach, grasp, or manipulate objects, vision of the arm provides information that is richer than a cursor, including cues about the limb’s shape, posture, orientation, and identity (whether the seen arm is a right or left arm). These cues may influence how movements are planned and controlled, yet they are often absent from many traditional motor control paradigms. As cognitive neuroscience research increasingly moves toward more naturalistic stimuli and tasks (Gibson 1997; Snow & Culham 2021), the use of more ecologically valid stimuli may enable a better understanding of how action is controlled in everyday life.

A growing body of behavioural evidence suggests that vision of the arm can influence how movements are controlled. Early work demonstrated that vision of the moving limb improves the control of aiming movements (Carlton 1981; Spijkers & Spellerberg 1995), and that visual feedback of the hand during reach-to-grasp actions can alter movement execution, including the timing of grip closure as participants prepare to grasp an object (Connolly & Goodale 1999; Rand et al. 2007). Converging evidence suggests that the visual form of the hand itself and the richness of information it provides may also influence action. Reaching and grasping performance can depend on the richness of visual hand feedback, with more detailed hand information producing better performance than reduced or simplified feedback (Brand et al. 2016; Sivakumar et al. 2021). People may also learn movements more effectively when they see an arm compared with other effectors, such as a tool (Shin et al. 2022). These findings suggest that the visual appearance of the arm may be integrated into internal representations of the acting limb, including moment-to-moment estimates of where the limb is, how it is positioned, and how it is moving. In this way, seeing an arm may support action not only by providing spatial feedback about where the limb is, but also by engaging body-specific representations that influence how movements are planned, monitored, and corrected.

Neuroimaging provides further evidence that vision of the arm may be important for action, showing that the visual system contains regions that respond selectively to the human body and hands. Human neuroimaging studies have identified brain areas that respond selectively to images of the body, including the extrastriate body area (EBA; Downing et al. 2001) and regions of the lateral occipitotemporal cortex (LOTC) that respond selectively to body parts such as hands (Bracci et al. 2010; Orlov et al. 2010). These hand-selective visual regions may represent action-relevant information, including information about how objects can be grasped (Freud et al. 2018; Knights et al. 2021). Although much of this work has focused on visual perception of the body, other evidence suggests that these regions may also contribute to action by generating expectations about visual feedback from the hand and comparing those expectations with actual sensory feedback during movement (Astafiev et al. 2004; van den Heiligenberg et al. 2018; Zimmermann et al. 2016).

Perhaps one of the most powerful ways to probe the influence of vision on the motor system is to create a mismatch between the limb that is seen and the proprioceptive signals arising from the limb that is actually moving. Mirror feedback uses the reflection of one limb to create the impression that the opposite limb is moving, and is thought to provide visual input that can engage sensorimotor areas associated with the reflected limb (Ramachandran & Altschuler 2009). Consistent with this idea, studies have shown that viewing a mirrored limb can modulate activity or excitability in primary motor cortex (M1) contralateral to the visually presented limb, even when that limb is not the one physically performing the action (Garry et al. 2005; Senna et al. 2015; Tominaga et al. 2011; Touzalin-Chrétien et al. 2009). The behavioural relevance of this visual influence is further supported by mirror therapy, which has been used in rehabilitation contexts such as stroke recovery and phantom limb pain (Ramachandran et al. 1995; Ramachandran & Altschuler 2009; Thieme et al. 2018). Together, this work suggests that seeing a body part move can have consequences for motor-related processing, experience, and behaviour, even when the seen limb does not directly correspond to the moving limb.

Although these findings suggest that vision of the arm can influence motor control, they do not fully isolate the contribution of the arm’s visual identity from the contribution of the physical arm performing the action in online motor control. One reason this issue remains difficult to study is that, in the real world, visual and motor properties of arm actions are typically congruent. For example, when an individual performs an action with their right arm, they typically see that same arm acting in space, resulting in congruent visual and motor information. This natural coupling makes it difficult to determine whether motor performance depends primarily on the arm that is physically acting, whether it is also influenced by the arm that is visually presented, or whether performance depends on the correspondence between the two.

Virtual reality (VR) offers a unique opportunity to study vision and action because it allows the visual representation of the body to be manipulated in ways otherwise impossible in the real world (Kilteni et al. 2012; Sanchez-Vives & Slater 2005; Slater et al. 2010). The use of virtual avatars in video games and VR therefore holds strong potential for addressing novel questions in sensorimotor control. In contrast to traditional experimental setups, VR enables researchers to systematically violate natural sensorimotor contingencies while preserving goal-directed behaviour (Mangalam et al. 2023). This makes VR particularly well suited for isolating the contribution of specific sensory information to sensorimotor control.

In VR, a participant can perform movements with one arm while receiving visual feedback from a different virtual limb, such as when the right arm controls a virtual left arm. By creating congruent and incongruent pairings between the acting arm and the seen arm, VR makes it possible to test whether motor control is solely shaped by the acting limb or also by visual information about the seen limb (Culham 2016; Ossmy & Mukamel 2016).

Studies of virtual bodies and virtual arms suggest that the way the body is visually presented in VR can influence both the embodiment of an avatar and action. One aspect of embodiment is ownership, or the feeling that the virtual body belongs to oneself. Ownership can extend to virtual arms that are anatomically implausible, such as switched left and right arms, as long as the virtual limbs move in synchrony with the participant’s own movements (Yizhar et al. 2021). In another line of research, when the perceived position of a virtual hand is spatially displaced, reaching movements are biased toward the location of the virtual hand (Ambron et al. 2020).

Indeed, neuroimaging also suggests that virtual video games can engage neural mechanisms similar to those involved in real actions (Leppala 2022). Unsurprisingly, activation in motor (M1) and somatosensory (S1) cortices was contralateral to the physical arm used to control a gamepad. More interestingly, visuomotor brain regions in parietofrontal cortex and the ventral visual stream depended more strongly on the appearance of the virtual arm seen performing an action. Furthermore, in right-handed participants, greater contralateral-hemisphere activation was observed when they controlled a virtual left arm compared with a virtual right arm, suggesting that seeing the less dextrous left arm may require additional neural processing. Behavioural performance was affected by the virtual arm but not the controlling arm. However, because of the physical constraints of the fMRI scanner, participants controlled the movements using a video game controller rather than performing actual movements that would provide rich proprioceptive information from the controlling arm. This leaves open the question of whether the visual identity of the arm can influence motor performance during more naturalistic tasks, such as actual reaching.

The present study used VR to dissociate the acting arm from the seen arm during action. By independently manipulating the physical arm used to perform the movement and the virtual arm displayed in VR, this study allowed for the examination of whether visual information about the body influences online motor control beyond simply indicating the position of the moving hand.

Two features were systematically manipulated, *Controlling Arm* and *Virtual Arm*, in a centre-out reaching task, in which participants reached from a central home target to one of eight peripheral targets spaced 45° apart (Fig. 1b). The four experimental conditions differed in the physical arm used to control the reaching movement (Controlling Right or Left) and the virtual arm displayed in the headset (Virtual Right or Left) (Fig. 1c). In the VR-CR and VL-CL conditions, participants saw a virtual arm that matched the controlling arm (congruent conditions). In contrast, in the VR-CL and VL-CR conditions, the virtual arm did not match the arm performing the movement (incongruent conditions). Examining performance across the eight targets allowed for the assessment of whether the effects of *Controlling Arm* and *Virtual Arm* depended on movement direction. Therefore, varying *Controlling Arm*, *Virtual Arm*, and *Target* resulted in a 2 × 2 × 8 factorial design.

**Fig. 1.**
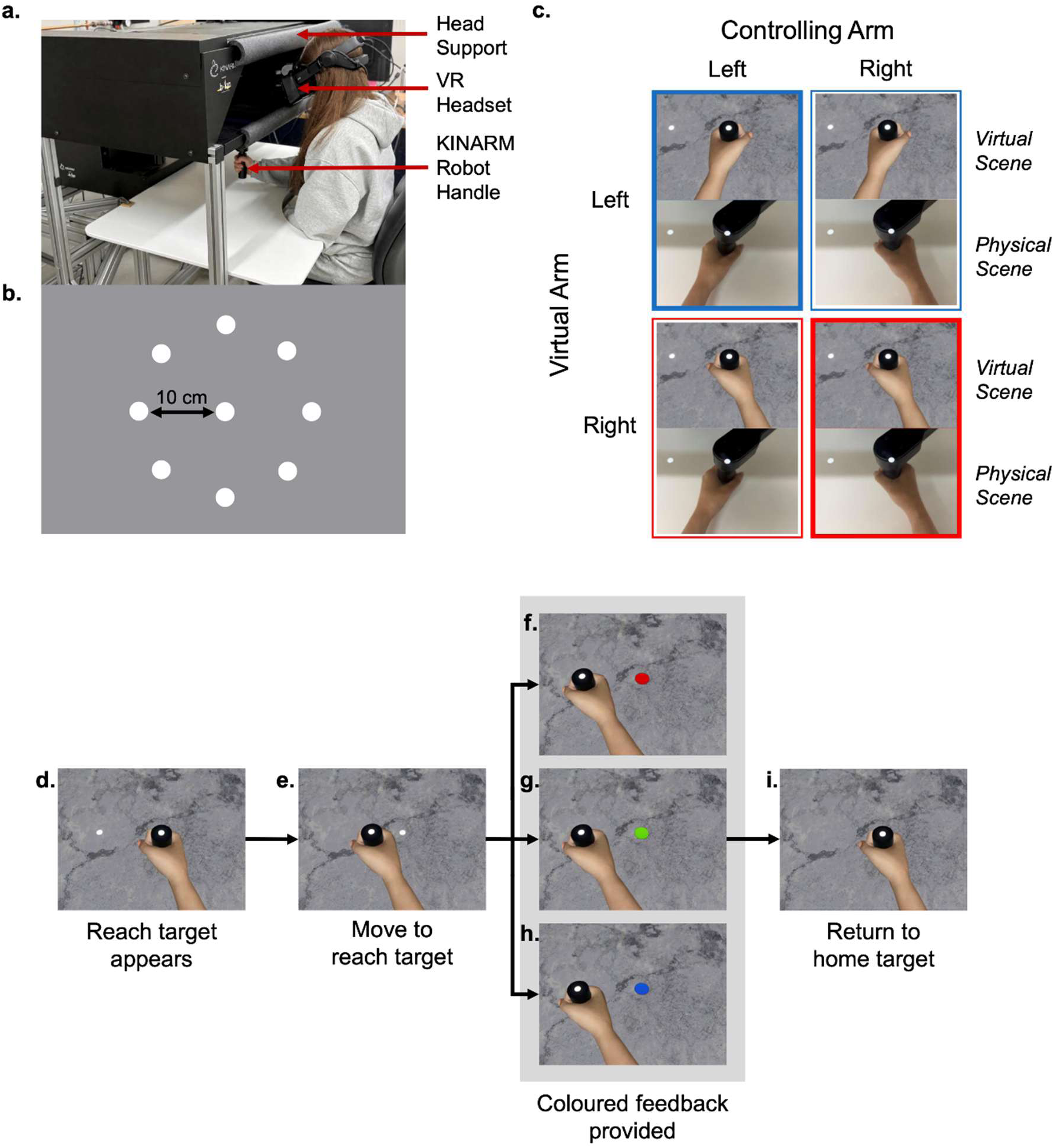
Experimental setup and trial design. (**a**) KINARM robot and VR setup. Participants were seated at the robot while wearing a VR headset. (**b**) Schematic of the centre-out reaching task. Participants reached from a central home target to one of eight possible reach targets. Hand position was indicated by a cursor centred on the KINARM handle. (**c**) Experimental conditions in the 2 × 2 design, which manipulated the controlling arm and the virtual arm displayed in VR. (**d –i**) Trial sequence design. (**d**) A reach target appeared. (**e**) Participants reached toward the target; upon leaving the home target, the reach target disappeared. Participants then received colour-coded feedback. (**f**) Red indicated that the reach was completed too quickly. (**g**) Green indicated that the reach was completed within the desired time window of 425 ± 50 ms. (**h**) Blue indicated that the reach was completed too slowly. (**i**) The robot returned the participant’s hand to the home target before the next trial

This design allowed the effects on motor performance of the seen arm to be separated from those of the acting arm. If there were no effect of the virtual arm, the results would show only an effect of the controlling arm, indicating that visual information about the identity of the moving limb does not meaningfully alter motor performance. If congruent conditions produced better performance than incongruent conditions, this could suggest that sensorimotor conflict or disruptions to the embodiment of the virtual avatar may have impacted movement. Alternatively, performance may depend on the identity of the virtual arm itself, such that seeing a right arm improves performance. In right-handed participants, the visual identity of the arm may carry different learned associations about action capability. The right arm is typically more practiced and dextrous, whereas the left arm is often used less during everyday actions. As a result, seeing a right arm may provide visual information that is more consistent with participants’ typical motor experience, potentially engaging neural mechanisms associated with the more dextrous right limb. Specifically, because all participants were right-handed, comparisons between conditions in which they saw their dominant right arm versus their non-dominant left arm tested whether the visual presentation of a more dextrous limb would enhance motor performance. This would provide evidence that vision of the dominant right arm can benefit precise motor control. It was predicted that seeing the right arm would improve performance, whereas seeing the left arm would reduce performance, even when the controlling arm remained the same.

## METHODS

### Participants

Data were analyzed from 36 participants (18 male, 18 female; mean age 21), recruited through Western University’s psychology research pool or through advertisements on campus. All participants were right-handed, as measured by the Edinburgh Handedness Inventory (Oldfield 1971), with strong right-hand preference, as indicated by a score of at least +40. All participants reported normal or corrected-to-normal visual acuity. To ensure that participants could perceive stereoscopic depth cues in the VR environment and to reduce the likelihood that task performance was confounded by deficits in stereo vision, stereoacuity was assessed using the Randot Preschool Stereoacuity Test (Stereo Optical Company Inc.). All participants included in the study could successfully identify disparities of 40 arcsec or better and had no history of strabismus or amblyopia. Prior to testing, all participants provided informed consent. Following completion of the study, participants received either financial compensation ($12/hour) or course credit for their time. All study procedures were approved by Western University’s Health Sciences Research Ethics Board in accordance with the standards of the 1964 Declaration of Helsinki.

### Overview

Participants were seated at a KINARM robotic manipulandum while wearing a VR headset that displayed a virtual environment aligned with the physical workspace of the robot (Fig. 1a).

Participants grasped the robot handle with either their right or left hand and performed centre-out reaching movements from a home target to one of eight possible reach targets (Fig. 1b). The design of the experiment allowed for the independent manipulation of the arm used to control the movement (right or left) and the virtual arm displayed in the headset (right or left; Fig. 1c). The structure of each trial is shown in Fig. 1d–i. Participants were instructed to move in a straight line to the reach target, each located 10 cm from the home position. The target disappeared upon movement onset such that endpoint accuracy could not be optimized using visual feedback.

Movement time was measured from the moment participants left the home target to the moment they arrived at the reach target. Participants were instructed to complete each reach within the desired time window of 425 ± 50 ms to ensure that accuracy measures of performance were not driven by large differences in the temporal structure of the reaches. Participants then received colour-coded feedback about their movement time. After each reach, the robot returned the participant’s hand to the home target before the next trial.

### Experimental Setup

Participants wore a Valve Index VR headset (Valve Corporation, Bellevue, WA, USA) and grasped the handle of a KINARM two-joint robotic manipulandum (KINARM Laboratories, Kingston, ON, Canada) with either their left or right arm to perform planar reaching movements to eight visual targets. The Valve Index was selected because it offers a relatively wide field of view compared with many modern VR headsets. The headset uses a dual-display system, with one screen per eye, and provides a field of view of 108° horizontally by 104° vertically, a per-eye resolution of 1440 × 1600 pixels, and a refresh rate of 120 Hz. The distance between the screens could be adjusted on the headset to an interpupillary distance (IPD) range of 58–70 mm. The KINARM had a refresh rate of 1000 Hz. Foam head supports were placed on the exterior of the robot to support participants’ heads and prevent neck strain from the weight of the VR headset during the experiment (Fig. 1a).

The VR setup was calibrated to match the physical space of the KINARM robot. To ensure that the VR setup was aligned to the physical workspace of the KINARM, the Open Sound Control (OSC) message protocol was used to transmit information from the KINARM experiment computer to the outboard VR computer running the Unity application. An OSC server embedded within the Unity application continuously listened for incoming messages from the KINARM, which carried both cursor position coordinates and event codes specifying which targets to render. Since the KINARM Simulink library does not natively include blocks for sending OSC messages, custom MATLAB script blocks were developed to fulfill this role. These blocks collected data from various components of the KINARM experiment, assembled it into properly formatted OSC messages, and transmitted them via the UDP Simulink block. The Unity OSC server was implemented using an open-source library developed by Thomas Fredericks (https://github.com/thomasfredericks/UnityOSC). Network connectivity between the two systems was established by connecting the VR computer to a Netgear GS110TP 8-Port network switch via a CAT6A ethernet cable, the same switch to which the KINARM robot and the experiment computers were also connected. This configuration allowed the VR computer and KINARM experiment computer to communicate directly with one another over a shared local network to ensure that manipulations completed in the physical workspace of the KINARM were reflected in VR. To ensure a stable and consistent connection, the VR computer was assigned a static IP address within the permitted range specified by the KINARM manual. The time delay between the KINARM handle position and the displayed virtual hand position was ∼9–10 ms. Graphics for this experiment were created and run in Unity, while the reaching task was created using MATLAB 2015a and run on the KINARM through Dexterit-E 2019.

Avatar models for the body and arms were created to be customizable to individual participants. Models were generated using Character Creator 3, based on default body templates modified with morphological options from the Ultimate Morphs add-on pack. Participants were asked to stand before the onset of the experiment so their height could be measured relative to the headset position. To better match the avatar’s size to that of a participant, a scaling ratio was calculated by dividing the participant’s measured height by the default height of the selected avatar model. This ratio was then applied uniformly across the x, y, and z axes of the avatar’s transform component, scaling the digital figure proportionally to better approximate the participant’s physical dimensions. For example, if the selected avatar had a default height of 180 cm and the participant’s height was 171 cm, the resulting ratio of 0.95 would be applied to reduce the avatar’s size accordingly. The body type and skin tone of the virtual arm could be adjusted to participants’ preferences to provide a more accurate representation of their own body.

Participants could choose one of two avatar bodies, with one being more muscular than the other. The posture of the avatar was adjusted to match the typical posture that participants would have during a standard KINARM reaching task. Participants were asked to limit their wrist movement during the experiment to ensure consistent posture with the avatar throughout the entire session.

Participants saw either a virtual left or right arm. The virtual hand grasped a 10-cm-high handle with a circular white cursor overlaid upon the top centre position of the handle that indicated its location in space. Targets were displayed through the VR headset and appeared horizontally on the same plane as the top of the handle. Targets were arranged equally around the circumference of a 10-cm-radius circle, with a home target at its centre (Fig. 1b). The virtual table below the targets was generated with a marble texture to provide rich visual cues about surface depth and orientation relative to the surface plane.

### Experimental Procedure

Screening and calibration procedures were completed prior to the experimental task. Interpupillary distance was measured using a pupilometer and entered into the VR headset to ensure accurate 3D depth perception and alignment of virtual objects with the participant’s real-world view. Before beginning the experimental task, participants completed 10 practice trials with the right arm and 10 practice trials with the left arm without the VR headset to get them accustomed to the reaching task.

On a given trial, participants began with the cursor representing the position of the KINARM robot handle in the home target, until prompted by the appearance of one of eight possible reach targets that were located 10 cm away from the home target. The reach target disappeared as soon as the cursor left the home target such that endpoint accuracy could not be optimized by continuous visual feedback. Participants were instructed to make their movements from the home target to the centre of where they saw the reach target appear in as straight a line as possible and to do so within the desired time window (425 ± 50 ms). After 500 ms of no movement, the trial ended and the robot moved the participant’s passive arm so that the cursor was placed back at the home position (Fig. 1d–i).

To help ensure that differences in spatial measures of performance were not driven by large individual differences in strategy (e.g., prioritizing accuracy vs. speed) and to keep the task sufficiently demanding, participants received coloured feedback after each reach to indicate whether the reach was completed too quickly (home target turned red), too slowly (home target turned blue), or within the desired window (425 ± 50 milliseconds; home target turned green). These time constraints included only the movement time of the participant. In other words, the coloured feedback reflected the amount of time elapsed between leaving the home target and ending the reach for a given trial.

The experimental procedure lasted approximately 50 minutes. The experiment was divided into four blocks of 120 trials, with each block taking approximately 7–8 minutes. Each block varied in the physical arm used to perform the task (right or left) and the virtual arm that was displayed to participants (right or left). In each block, each of the eight reach targets appeared fifteen times per block in random order. The order in which the blocks were presented to participants was counterbalanced across participants.

### Data Analysis

The kinematic data from participants’ reaches were sampled at 1000 Hz. The data were then low-pass filtered using a second-order Butterworth filter with a cutoff frequency of 10 Hz implemented in MATLAB.

Quality assurance was performed by first plotting each participant’s trajectory data and examining the plots (Fig. 2a). This was done to ensure that all participants had correctly followed the instructions of the reaching task, such as reaching to the correct target during each trial.

**Fig. 2.**
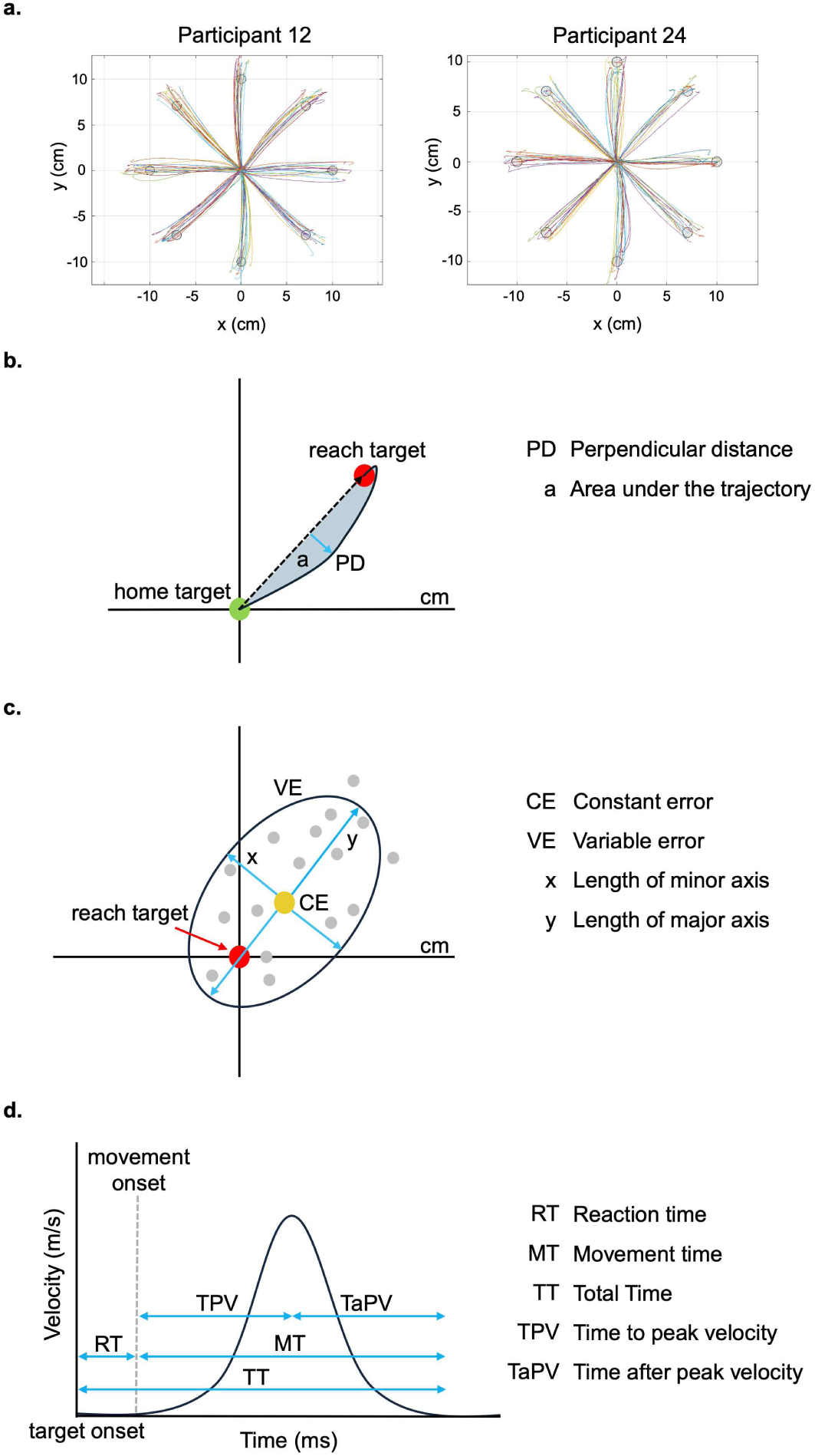
Trajectory plots and schematics of accuracy and timing variables. (**a**) As part of data quality assurance, trajectory data were plotted for each participant and condition (VR-CR, VR-CL, VL-CR, VL-CL) and examined. (**b**) Schematic of curvature error. The maximum perpendicular distance (PD), represented as a blue arrow, indicates the greatest unsigned deviation from a straight-line path. Area under the trajectory, represented as the shaded blue area, indicates the cumulative area between the observed hand path and a straight-line path. (**c**) Schematic of endpoint error. The constant error (CE), represented as the yellow circle, indicates the average location of endpoints relative to the reach target. Variable error (VE) was summarized with an ellipse that encompassed 95% of the trialwise reach endpoint locations. The lengths of the minor (x) and major (y) axes describe the shape of the variable error ellipse. (**d**) Shown is a velocity-time graph that represents the movement of the participant’s hand throughout a trial and depicts how each timing variable is measured

Reaches were excluded in cases in which a participant sped up and slowed down multiple times between leaving the home target and reaching the reach target, as determined by the presence of two or more acceleration phases separated by a deceleration phase (Mangos et al. 2024). Fewer than 5% of trials were excluded based on this criterion.

Before analysis, trajectories were rotated so that each target direction was aligned to a common reference axis, with the reach target positioned directly above the home target. This allowed trajectory measures to be compared within a shared coordinate frame, regardless of the original target location. Statistical analyses were conducted with the statistical software JASP (version 0.19.3) and Jamovi (version 2.7.33).

### Dependent Variables

To assess performance across the four experimental conditions, measures of movement accuracy and timing were examined (Fig. 2).

Maximum perpendicular distance was defined as the greatest unsigned deviation of the hand path from the ideal straight-line path between the home target and reach target. Area under the trajectory was calculated as the cumulative area between the observed hand path and the ideal straight-line path, providing an overall measure of trajectory deviation across the movement, making it sensitive to sustained curvature and late corrections toward the target (Fig. 2b). As shown in Fig. 2a, several trajectories exhibited a hook-like deviation near movement termination. Compared with maximum perpendicular distance, area under the trajectory is better able to capture these late corrections because it reflects deviations across the full movement path.

Constant error was computed separately along the reach axis and perpendicular axis by averaging trialwise endpoint errors within each participant and condition, with the reach axis reflecting error in the direction of the reach and the perpendicular axis reflecting error perpendicular to the reach path. This measure captured systematic directional bias in endpoint location for each target. Variable error was quantified by constructing endpoint variability ellipses for each participant and condition from the covariance matrix of reach endpoint locations. Individual covariance matrices were then averaged across participants to derive a group-averaged 95% confidence interval (CI) ellipse for each condition and target. To assess whether the variable error ellipses differed across conditions, statistical analyses were performed on the x- and y-coordinates of the ellipse centres and the lengths of the major and minor axes (Fig. 2c).

To assess movement timing across conditions, the two-dimensional position of the cursor over time was analyzed to derive reaction time, movement time, and total time. Movement time was also divided into time to peak velocity and time after peak velocity measured in milliseconds (Fig. 2d). Reaction time was defined as the interval between the reach target appearing and movement onset. Movement time was defined as the interval between movement onset and the end of the movement. Summing together reaction time and movement time, total time was derived. Time to peak velocity was defined as the interval between movement onset and the moment of peak tangential velocity. Finally, time after peak velocity was defined as the interval between the moment of peak tangential velocity and movement end and was used as an index of the later phase of movement, including online correction and deceleration processes. For each participant, median values across trials were extracted for the timing variables because medians are more resistant to skewed distributions, without requiring arbitrary decisions about outlier removal.

For each dependent variable, a 2 (controlling arm) × 2 (virtual arm) × 8 (target location) ANOVA was performed. Although some variables showed interactions of target location with *Controlling Arm* and/or *Virtual Arm*, these interactions were generally explained by effects in the same direction (e.g., better performance for CR than CL) that differed in magnitude but not direction. As such, to simplify the main data presentation, target location was collapsed and the results were focused on the main effects and interactions for *Controlling Arm* and *Virtual Arm.* Interactions of these variables with *Target* are reported in the Supplementary Information. A Bonferroni correction was applied to *post hoc* tests.

## RESULTS

### Accuracy

#### Curvature Error

Curvature error was assessed using two complementary measures: maximum perpendicular distance, which captures the largest single deviation from a straight reach path, and area under the trajectory, which captures cumulative deviation across the full reach.

Maximum perpendicular distance was examined first. As shown in Fig. 3a, trajectories were straightest when participants both controlled and viewed a virtual right arm (VR-CR; solid red circle) compared with the other three conditions (VR-CL, VL-CR, and VL-CL). This pattern was supported by a three-factor ANOVA, which revealed an interaction between *Controlling Arm* and *Virtual Arm* [*F*(1, 35) = 6.4, *p* < 0.05, *η^2^_p_* = 0.16]. This result suggests that trajectory curvature was not determined solely by the physical arm used to perform the movement. Instead, the visual identity of the arm also influenced movement control, with the greatest performance emerging when the right controlling arm was paired with a visually congruent virtual right arm. In addition to the interaction, the ANOVA also revealed main effects of *Controlling Arm* [*F*(1, 35) = 5.7, *p* < 0.05, *η^2^_p_* = 0.14], *Virtual Arm* [*F*(1, 35) = 11.0, *p* < 0.01, *η^2^_p_* = 0.24], and *Target* [*F*(7, 245) = 10.7, *p* < 0.001, *η^2^_p_* = 0.23]. *Target* also interacted with *Controlling Arm* [*F*(7, 245) = 7.0, *p* < 0.001, *η^2^_p_* = 0.17] and *Virtual Arm* [*F*(7, 245) = 2.5, *p* < 0.05, *η^2^_p_* = 0.07]. However, these interactions appeared to reflect quantitative rather than qualitative differences and were generally explained by effects in the same direction that differed in magnitude but not direction (e.g., less curvature error for CR than CL). These target-specific effects are presented in Supplementary Fig. 1a for completeness.

**Fig. 3.**
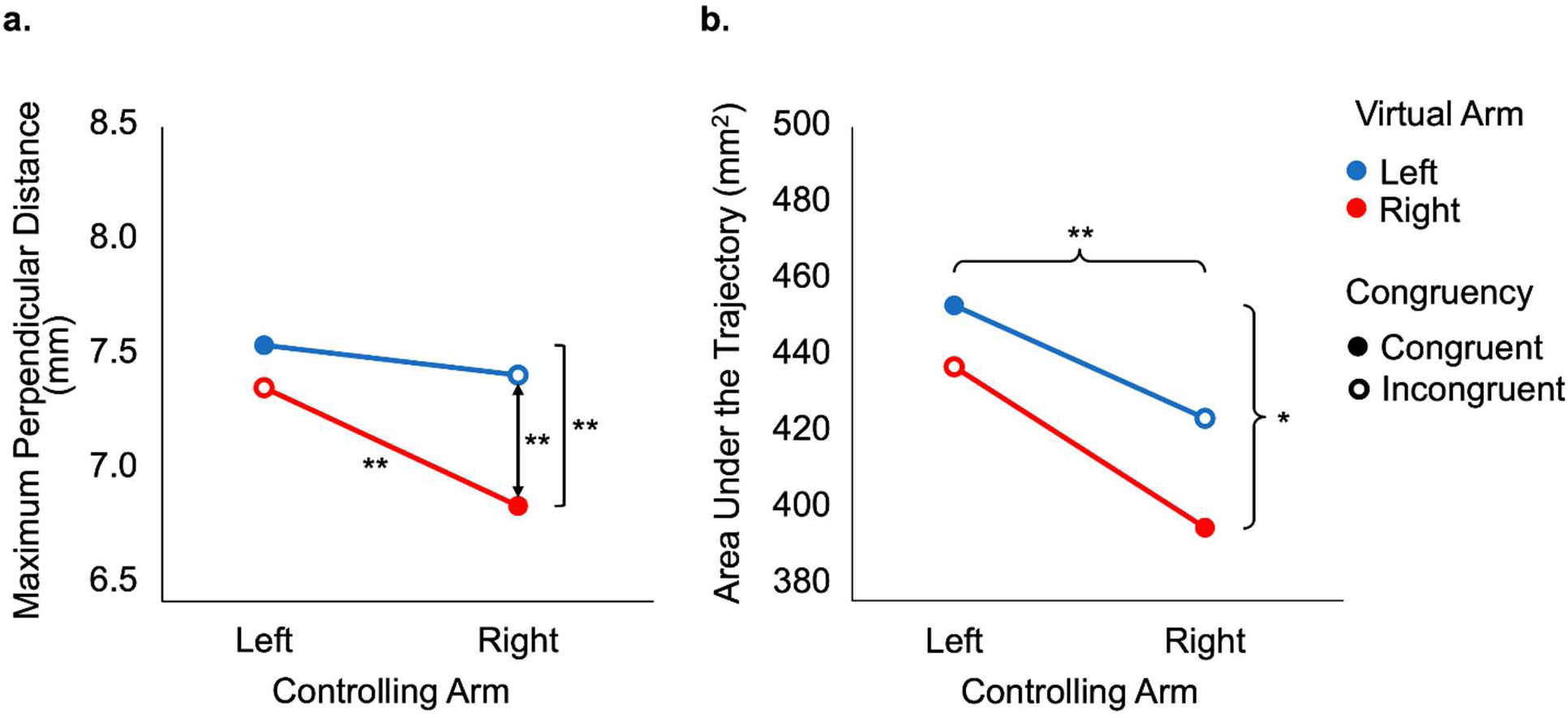
Curvature error across conditions. Main effects are shown with braces, and interaction effects are shown with brackets, arrows and stars above the lines. \**p* < 0.05, \*\**p* < 0.01, \*\*\**p* < 0.001 after *post hoc* analysis. (**a**) Interaction between *Controlling Arm* and *Virtual Arm* on maximum perpendicular distance. Participants had straighter reaches when they both saw and controlled a virtual right arm compared with the other three conditions. (**b**) Main effects of *Controlling Arm* and *Virtual Arm* on area under the trajectory. Participants’ reaches exhibited less curvature when they saw a virtual right arm compared with a virtual left arm within each controlling arm

We next examined area under the trajectory, which captures aspects of trajectory shape that may not be reflected by maximum perpendicular distance, including late corrective movements that can produce a hook-like shape near the end of the reach. As shown in Fig. 3b, trajectories were straighter when participants controlled the movement with their right arm compared with their left arm. Interestingly, trajectories also showed less overall curvature when participants viewed a virtual right arm, regardless of which arm controlled the movement. This observation was supported by a three-factor ANOVA, which revealed main effects of *Controlling Arm* [*F*(1, 35) = 10.2, *p* < 0.01, *η^2^_p_* = 0.23] and *Virtual Arm* [*F*(1, 35) = 4.5, *p* < 0.05, *η^2^_p_* = 0.11]. This result extends the maximum perpendicular distance finding by showing that viewing a virtual right arm reduced curvature across the entire reach. Notably, this effect was also present when participants controlled the movement with their left arm, suggesting that the visual appearance of the arm could improve trajectory control even when it was dissociated from the acting limb. The ANOVA also revealed a main effect of *Target* [*F*(7, 245) = 6.2, *p* < 0.001, *η^2^_p_* = 0.15] and an interaction between *Controlling Arm* and *Target* [*F*(7, 245) = 5.0, *p* < 0.001, *η^2^_p_* = 0.13]. As with maximum perpendicular distance, the target-related effects were generally driven by the same overall pattern across targets, with the effects of *Controlling Arm* varying in magnitude rather than reversing direction (Supplementary Fig. 1b).

#### Endpoint Error

Next, endpoint error was evaluated to determine how accurately participants reached to the reach target locations. To assess endpoint error, average 95% CI ellipses centred on the average x, y locations of where participants ended their reaches for each condition were generated (Supplementary Fig. 2). Across all conditions, participants tended to overshoot along the reach axis. One possible explanation is that the reach target disappeared at movement onset, requiring participants to guide their movements based on a remembered target location rather than continuous visual information. Overall, participants showed less average error in the reach direction when controlling the movement with their right arm compared with their left arm. Detailed analyses of constant error are depicted in Supplementary Figs. 3–5.

The variable error of the endpoints was assessed using the length of the major axis (along the reach direction) and minor axis (perpendicular to the reach direction) of the variable error ellipses. Variability along the reach direction was generally lower for controlling right-arm reaches than left-arm reaches. There was also a statistical trend (*p* < 0.1) suggesting that viewing a virtual right arm may have reduced endpoint variability along both axes compared with viewing a virtual left arm within each controlling arm. Detailed analyses of variable error are depicted in Supplementary Figs. 6–7.

#### Timing

Lastly, timing variables were assessed to examine differences in the timing of participants’ reaches. Reaction times were unexpectedly longer for right-arm reaches than left-arm reaches, which may suggest that participants took slightly longer to prepare movements with the right arm. One interpretation is that right-arm reaches involved greater preparation for better-controlled movements. In contrast, movement times and times after peak velocity were shorter for right-arm reaches than left-arm reaches, suggesting that once initiated, right-arm movements were completed more quickly. Total time did not appear to show any main effects, indicating that differences in reaction time and movement execution may have offset one another. Time to peak velocity also did not show any main effects; however, an interaction between *Controlling Arm* and *Target* revealed some target-specific differences. Generally, each arm appeared to take longer to reach peak velocity for targets on its own side of space. Overall, because these timing measures depended on the controlling arm rather than the virtual arm, the effects of virtual arm identity on accuracy were unlikely to be explained by a speed-accuracy trade-off. All main effects and interactions for timing are depicted in Supplementary Figs. 8–10 for completeness.

## DISCUSSION

### Summary of Results

Our results show that reaching performance was influenced, as expected, by the controlling arm, but more interestingly, also by whether the virtual display depicted a right or left arm (VR or VL). Trajectories were straightest when participants viewed a virtual right arm, with the straightest movements occurring when the right controlling arm was paired with a virtual right arm. Thus, seeing the virtual right arm improved overall trajectory control even when participants controlled the less dextrous left arm. Conversely, viewing a virtual left arm appeared to reduce accuracy when participants controlled the right arm. Thus, the effect was not explained solely by the physical arm used to perform the movement or by simple congruency between the acting and seen arm. Instead, the appearance of the virtual arm influenced motor performance.

Reach endpoints showed less error when participants controlled the movement with their right arm compared with their left arm. However, there was also a trend toward lower variability across trials when participants saw a virtual right versus left arm, suggesting that visual arm identity may also influence how consistently movements are brought to the target. In contrast, timing measures were influenced only by the controlling arm, suggesting that the accuracy effects were not simply driven by a speed-accuracy trade-off. Instead, the results point to a dissociation between spatial and temporal aspects of performance: the identity of the virtual arm influenced accuracy measures together with the controlling arm, whereas timing measures were affected solely by the controlling arm.

Taken together, these results indicate that the appearance of the identity of the arm may play a more important role in motor control than previously assumed. The findings highlight the importance of considering body-related visual information in visuomotor studies and demonstrate the value of VR as a tool for studying how the seen body contributes to sensorimotor control.

### The Seen Arm Matters

Previous work has shown that seeing the arm can influence several aspects of visually guided action, including aiming (Carlton 1981; Spijkers & Spellerberg 1995), reaching and grasping objects (Connolly & Goodale 1999; Rand et al. 2007), and motor learning (Shin et al. 2022). The present study extends this work by showing that the influence of vision of the arm is not limited to whether the arm is visible or absent, or the richness of its appearance, but also the visual identity of the arm. The finding that trajectories were straighter when participants viewed a virtual right arm suggests that the identity of the seen arm can influence the control of movements.

One possible interpretation is that the visual identity of the limb biased the internal representation used to guide the reach toward one associated with the more practiced dominant arm. In right-handed individuals, the right arm is typically the more practiced and dextrous effector and may therefore be more strongly associated with skilled, precise action. This interpretation is consistent with evidence that the left hemisphere, which primarily controls the right side of the body, is specialized for general aspects of skilled action, praxis, and motor planning (Goodale 1988; Haaland et al. 2000; Johnson-Frey 2004; Perenin & Vighetto 1988). Handedness has also been shown to be associated with asymmetries in cortical hand representations. For example, the dominant right hand has been shown to have a larger somatosensory representation than the non-dominant left hand in the corresponding contralateral hemispheres (Sörös et al. 1999). Therefore, seeing a right arm may have cued sensorimotor processes normally associated with skilled, right limb action, even when the right arm was not the limb physically controlling the movement.

The present findings are consistent with previous neuroimaging evidence that the identity of the seen virtual arm can modulate brain activity within visuomotor regions (Leppala 2022). However, because this study used a gamepad and a 2D virtual display, it did not include direct proprioceptive or haptic feedback from a reaching limb, and the mapping between the physical movement and the virtual action differed from real-world reaching. The present findings expand on this work by showing a behavioural effect of virtual arm identity during real-world reaching. Other studies have similarly shown that the visual identity of the seen hand can modulate activity in action-related brain regions, including the anterior intraparietal cortex (Shmuelof & Zohary 2005, 2006). However, these studies used videos of isolated hands rather than fully intact visual limbs, such as in the present study. The current study extends these findings by showing that visual arm identity can influence behaviour in real-world reaching movements in a near-veridical 3D environment.

Previous work on ownership and agency has shown that anatomically implausible virtual arms, such as switched right and left arms, can still be experienced as part of the body when they move synchronously with the participant’s own movements (Yizhar et al. 2021). However, this study did not examine whether these manipulations affected movement kinematics. The present results suggest that even when virtual arms move in synchrony with the acting limb, their visual identity can still influence reaching behaviour.

Together, these findings suggest that the seen arm can shape sensorimotor processing even when it is dissociated from the acting arm. Building on this work, the present findings suggest that the visual identity of the arm can also influence real reaching behaviour. In right-handed participants, viewing a right arm may recruit sensorimotor networks associated with the dominant right limb, contributing to straighter movement trajectories. Future neuroimaging research, possibly using video games, may be able to further investigate this topic.

### Limitations

One limitation of the present study is that only right-handed participants were tested. As a result, it is difficult to determine whether the observed benefit of viewing a virtual right arm reflects a dominant/non-dominant arm effect or a right/left arm effect specific to right-handed individuals. Future work could examine whether similar effects are observed in left-handed participants, particularly because handedness effects may not be symmetrical across groups. For example, right-handed individuals have been shown to have a strong preference for using their dominant hand during precision grasping, whereas left-handed individuals do not always show the same degree of dominant-hand preference (Gonzalez et al. 2007).

The current state of VR limb rendering and tracking technology was another limitation of this study. The virtual arm was not fully matched to each participant’s real arm geometry and posture, despite efforts to make the alignment as accurate as possible. Wrist movements and elbow movements were not tracked. Instead, only the location of the KINARM robot handle was tracked, and participants were asked to align their posture with the virtual avatar throughout the experiment. Although this approach allowed the virtual arm to be presented consistently across participants, it may have introduced mismatches between the physical limb and the visual limb. Importantly, however, the virtual arm still influenced behaviour despite these imperfections. This is consistent with other work showing that artificial limbs can still influence body representation, as demonstrated by effects such as in the rubber hand illusion (Botvinick & Cohen 1998). Future systems with more complete upper-limb tracking could provide a more anatomically accurate visual representation of the acting limb and help determine whether reducing discrepancies between the seen and controlling arm changes the magnitude of the observed effects.

Another limitation is that although VR provides a powerful way to manipulate visual information of the body, it is not identical to real-world action. Factors such as display resolution, reduced field of view, and the vergence–accommodation conflict may influence performance in VR (Harris et al. 2019). The vergence–accommodation conflict occurs because the eyes converge to the simulated depth of the virtual object, while accommodation remains fixed on the physical display screen inside the VR headset. This creates a mismatch between two depth cues that are normally coupled in the real world. As a result, depth perception in VR may be less precise, which could have influenced participants’ perception of target or hand position.

### Future Directions

Future work could extend these findings in several ways. One direction is to examine applications relevant to rehabilitation for neurological conditions such as stroke and associated disorders of skilled action, including limb apraxia (Buxbaum et al. 2008), using approaches such as VR-based rehabilitation (Laver et al. 2025). Virtual displays of human bodies are increasingly used in interventions for motor impairment, including mirror therapy (Gebreheat et al. 2024).

Mirror therapy is thought to improve the function of the affected limb by creating the visual illusion that the impaired limb is moving successfully, which may increase engagement of action-observation, motor imagery, and sensorimotor networks in movement planning and control (Arya et al. 2015; Ramachandran & Altschuler 2009; Rossiter et al. 2015). The present findings raise the possibility that the visual identity of the limb shown in VR may be an important design feature in these interventions. For patient populations with upper-limb impairment, future work could test whether the visual identity of the virtual limb influences rehabilitation outcomes. In this approach, patients could move their unaffected limb while viewing a virtual version of their impaired limb performing the action successfully, allowing researchers to test whether this type of visual feedback strategy supports motor recovery of the impaired limb. This may be especially relevant for right-handed patients who lose function of their dominant right limb, as restoring successful movement of this limb may be particularly important for everyday action. Furthermore, VR could extend this approach by allowing patients to practice a wider range of movements and tasks without being restricted to a mirror setup.

Another direction is to examine whether the effects of virtual arm identity persist over time. The present study measured online motor performance, but it remains unknown whether repeated exposure to a particular virtual arm could produce longer-lasting changes in movement control. Future work could also examine whether learning of different actions transfers between the two arms depending on the virtual limb shown (Culham 2016; Ossmy & Mukamel 2016).

Intermanual transfer is a phenomenon that refers to the finding that practice with one arm can sometimes improve performance with the opposite, untrained arm, and has been shown in a multitude of different motor tasks (Malfait & Ostry 2004; Sainburg & Wang 2002; Stockinger et al. 2015). Testing whether virtual arm identity changes the amount or direction of intermanual transfer would determine whether the seen arm influences not only online movement control, but also the formation and sharing of newly acquired visuomotor mappings across limbs.

## Conclusions

The present study showed that reaching movements are shaped not only by the physical arm used to act, but also by the visual identity of the arm that is seen (right or left). By using VR to dissociate the acting arm from the seen arm, this study demonstrates that body-related visual information can influence motor control during reaching. The finding that viewing a virtual right arm improved the accuracy of movements in right-handed participants, regardless of the controlling arm, suggests that the appearance of the arm may contribute to internal representations used to plan, guide, and correct action. More broadly, these findings highlight that the visual environment, particularly vision of the body, is worth considering in the design of motor control studies.

## DATA AVAILABILITY

Data are available at: https://doi.org/10.5281/zenodo.23174377

## ACKNOWLEDGEMENTS

This research was supported by Discovery grants from the Natural Sciences and Engineering Research Council (NSERC) of Canada (04271-2022-RGPIN to J.C.C. and 04586-2025-RGPIN to P.L.G.). O.I. was supported by an NSERC Canada Graduate Scholarship – Master’s and an Ontario Graduate Scholarship. J.C.C. was supported by a Canada Research Chair in Immersive Neuroscience salary award. K.B. was supported in part by an operating grant from the Canadian Institutes of Health Research (PJT 190159). We thank William Naylen Sorrell for guidance with the KINARM system and Eva Deligiannis for helpful discussion and advice.

