## Supplemental Material for "Reaching movements in virtual reality depend on the seen avatar arm and not just the acting arm"

### Supplementary Information

#### Target Interactions for Curvature Error

A three-factor ANOVA was performed on the data for maximum perpendicular distance and area under the trajectory. These analyses identified an interaction between *Virtual Arm* and *Target* for maximum perpendicular distance, and interactions between *Controlling Arm* and *Target* for both variables. As shown in Supplementary Fig. 1, interactions appeared to reflect differences in magnitude, rather than the direction, of the effects across targets. Overall, error tended to be lower when participants either controlled or viewed a right arm compared with a left arm.

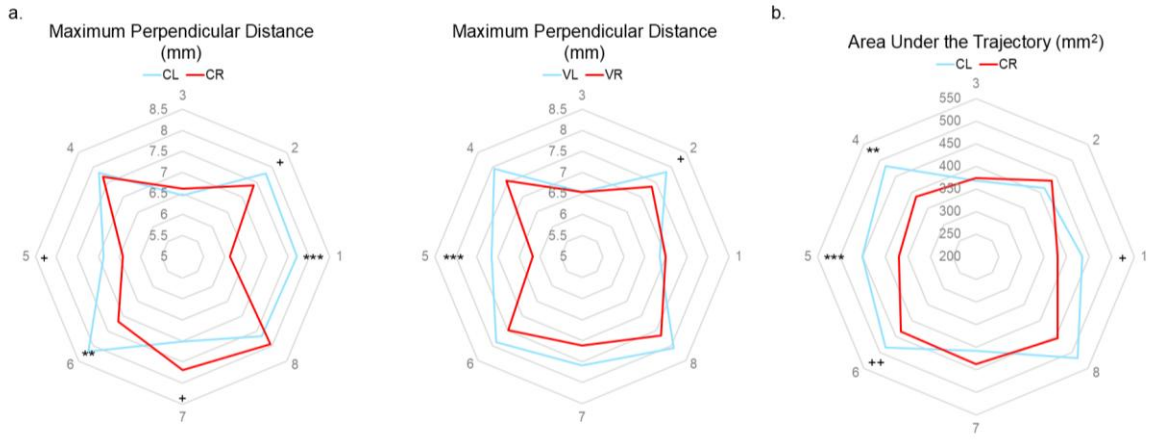


**Supplementary Fig. 1** *Controlling Arm/Virtual Arm* and *Target* interactions for curvature error. + $p_{uncorr} < 0.05$, ++ $p_{uncorr} < 0.01$. *$p < 0.05$, **$p < 0.01$, ***$p < 0.001$ after *post hoc* analysis. (**a**) *Controlling Arm* or *Virtual Arm* and *Target* interactions for maximum perpendicular distance. (**b**) *Controlling Arm* and *Target* interaction for area under the trajectory

#### Constant Error and Variable Error Ellipses

Endpoint variability was quantified by computing a covariance matrix from the reach endpoints for each participant and condition. These covariance matrices were then averaged across participants to generate a group-level 95% confidence ellipse for each condition and target. Visual inspection of Supplementary Fig. 2 suggests that across all conditions, participants tended to overshoot along the reach axis.

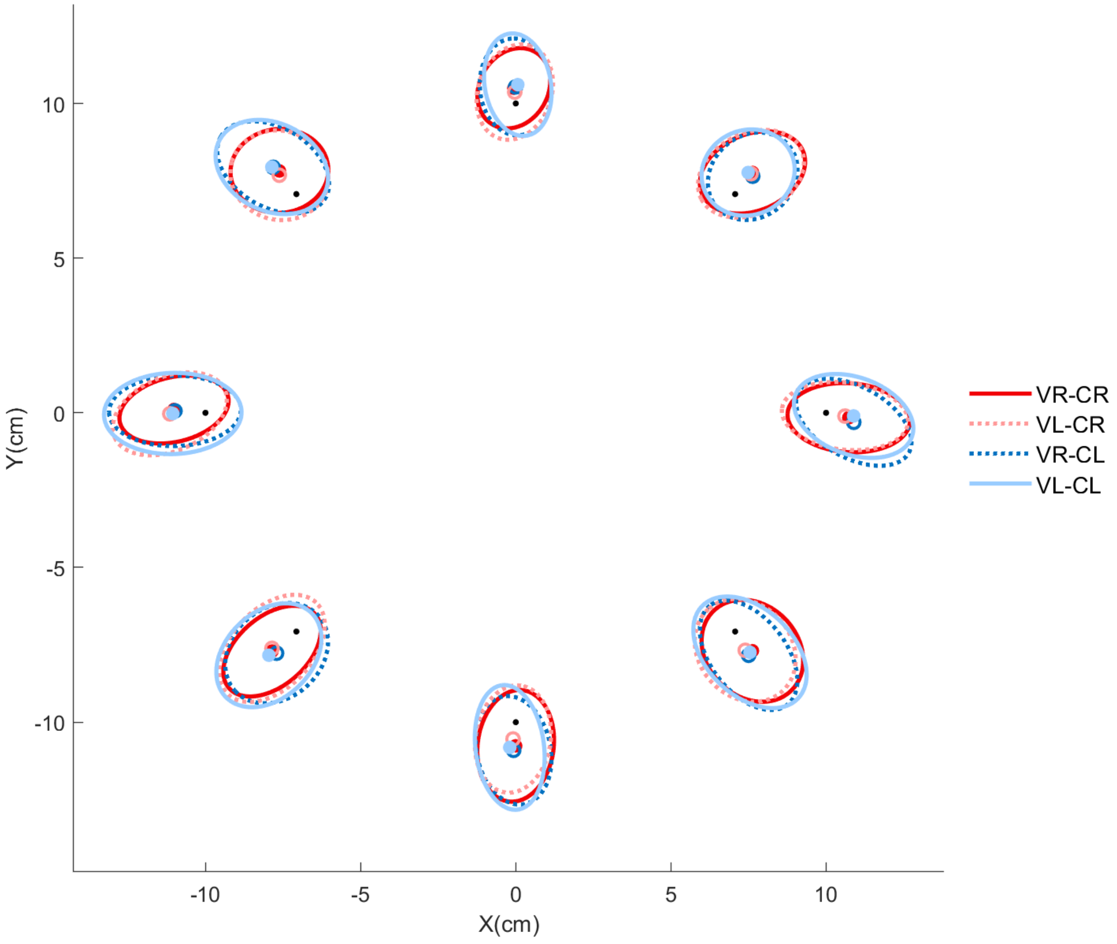


**Supplementary Fig. 2** Constant error and variable error ellipses. The black dot represents the location of each reach target. The coloured circles represent the x- and y-coordinates of the constant error for each target across participants. The ellipses represent the variable error for each target across participants

**Detailed Results on Constant Error**

Endpoint location in the reach direction, reflecting overshoot or undershoot, was assessed from the average y-position of the rotated endpoint data after trajectories were aligned so that the reach target was positioned directly above the home target. Participants overshot less in the reach direction when controlling the movement with their right arm compared with their left arm (Supplementary Fig. 3). This pattern was supported by a three-factor ANOVA, which revealed a main effect of *Controlling Arm* [$F(1, 35) = 4.7$, $p < 0.05$, $\eta^2_p = 0.12$]. The ANOVA also identified interactions between *Controlling Arm* and *Target*, $F(7, 245) = 3.52$, $p < 0.01$, $\eta^2_p =$ 0.09 and *Virtual Arm* and *Target*, $F(7, 245) = 3.65$, $p < 0.001$, $\eta^2_p = 0.09$, as well as a main effect of *Target,* $F(7, 245) = 8.92$, $p < 0.001$, $\eta^2_p = 0.20$. However, as shown in Supplementary Fig. 4, the effect of controlling arm only varied in magnitude and not direction whereby the controlling right arm showed less error than the controlling left arm.

A three-factor ANOVA on the data for the average x location identified a three-way interaction between *Controlling Arm*, *Virtual Arm*, and *Target* [$F(7, 245) = 3.0$, $p < 0.01$, $\eta^2_p = 0.08$], along with interactions between *Controlling Arm* and *Target* [$F(7, 245) = 2.3$, $p < 0.05$, $\eta^2_p = 0.06$], and *Virtual Arm* and *Target* [$F(7, 245) = 6.3$, $p < 0.001$, $\eta^2_p = 0.15$], as well as a main effect of *Target* [$F(7, 245) = 4.0$, $p < 0.001$, $\eta^2_p = 0.10$]. As depicted in Supplementary Fig. 5, there was limited evidence for reliable target-specific differences.

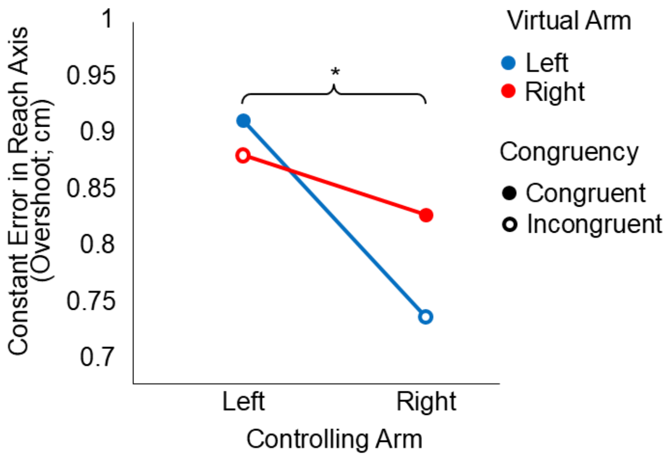


**Supplementary Fig. 3** Main effect of *Controlling Arm* for constant error in the reach axis (overshoot). *$p < 0.05$, **$p < 0.01$, ***$p < 0.001$. Main effects are shown with braces

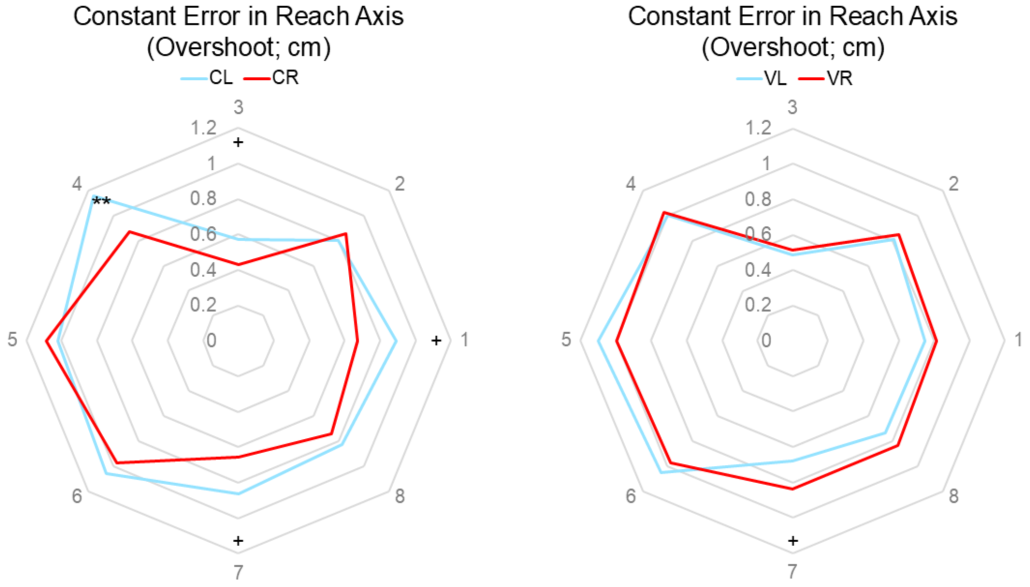


**Supplementary Fig. 4** *Controlling Arm*/*Virtual Arm* and *Target* interactions for constant error in the reach axis (overshoot). + $p_{uncorr} < 0.05$, ++ $p_{uncorr} < 0.01$. *$p < 0.05$, **$p < 0.01$, ***$p < 0.001$ after *post hoc* analysis

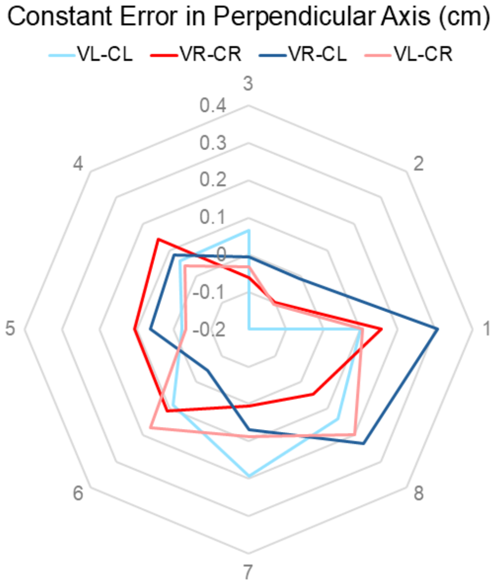


**Supplementary Fig. 5** *Controlling Arm*, *Virtual Arm* and *Target* interaction for constant error in the perpendicular axis (perpendicular to the reach). Positive values indicate error to the right of a straight-line trajectory, while negative values indicate error to the left of a straight-line trajectory. No comparisons were significant after *post hoc* analysis

**Detailed Results on Variable Error**

Endpoint variability was assessed using the length of the major (reach direction) and minor (perpendicular to the reach direction) axes of the variable error ellipses. Major axis lengths were shorter for right-arm reaches than left-arm reaches, suggesting less spread in endpoint positions along the reach direction when participants controlled the movement with their right arm (Supplementary Fig. 6a). This was supported by a three-factor ANOVA on the data for the major axis length, which revealed a main effect of *Controlling Arm* [$F(1, 35) = 5.3$, $p < 0.05$, $\eta^2_p = 0.13$]. There was also a statistical trend towards a main effect of *Virtual Arm* [$F(1, 35) = 4.1$, $p = 0.05$, $\eta^2_p = 0.11$], suggesting that endpoint variability along the major axis was generally lower when participants viewed a virtual right arm compared with a virtual left arm within each controlling arm. The ANOVA also revealed an interaction of *Controlling Arm* and *Target* [$F(7, 245) = 6.3$, $p < 0.001$, $\eta^2_p = 0.15$], and a main effect of *Target*, [$F(7, 245) = 18.3$, $p < 0.001$, $\eta^2_p = 0.34$]. Closer inspection of the interaction between *Controlling Arm* and *Target* showed that the left controlling arm generally produced more error than the right controlling arm, with the exception of the upper-right target (Supplementary Fig. 7a).

A similar pattern was observed for the minor axis length of the variable error ellipses. Endpoint variability along the minor axis tended to be lower when participants viewed a virtual right arm compared with a virtual left arm within each controlling arm (Supplementary Fig. 6b). In support of this, a three-factor ANOVA on the data for minor axis length revealed a statistical trend towards a main effect of *Virtual Arm* [$F(1, 35) = 3.6$, $p = 0.065$, $\eta^2_p = 0.09$]. The ANOVA also indicated an interaction of *Controlling Arm* and *Target* [$F(7, 245) = 3.7$, $p < 0.001$, $\eta^2_p = 0.09$], and a main effect of *Target* [$F(7, 245) = 12.0$, $p < 0.001$, $\eta^2_p = 0.26$]. As shown in Supplementary Fig. 7b, the effect of controlling arm only varied in magnitude and not direction whereby the controlling right arm showed less error than the controlling left arm.

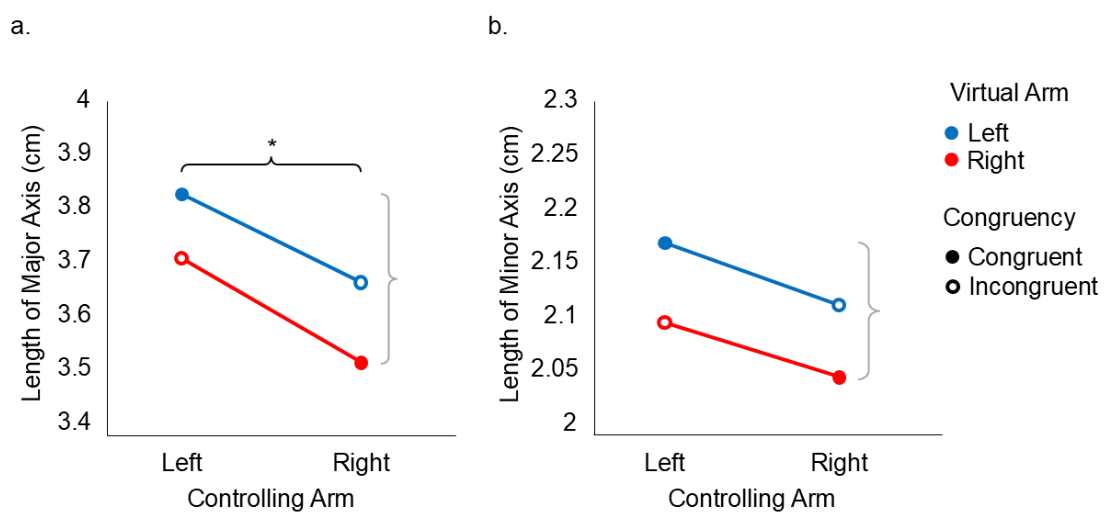


**Supplementary Fig. 6** Variable error across conditions. Main effects are shown with braces, with grey braces indicating trends. *$p < 0.05$, **$p < 0.01$, ***$p < 0.001$. (**a**) Length of the major axis (reach direction) of the variable error ellipse. A main effect of *Controlling Arm* and a trend towards a main effect of *Virtual Arm* ($p = 0.05$) were identified. (**b**) Length of the minor axis (perpendicular to reach direction) of the ellipses. A trend towards a main effect of *Virtual Arm* ($p = 0.065$) was identified

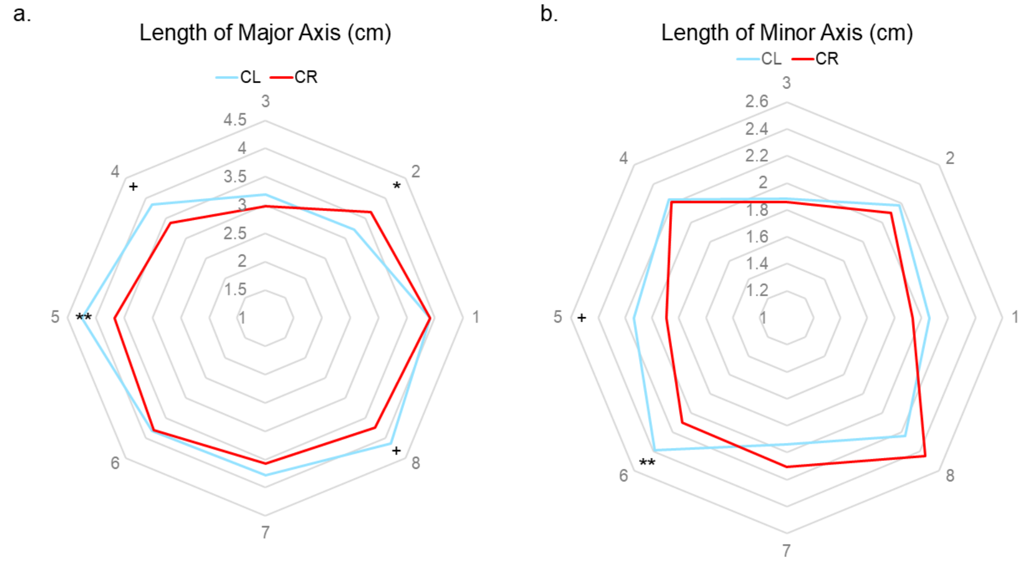


**Supplementary Fig. 7** *Controlling Arm* and *Target* interactions for the length of the major and minor axes of the ellipses. + $p_{uncorr} < 0.05$, ++ $p_{uncorr} < 0.01$. *$p < 0.05$, **$p < 0.01$, ***$p < 0.001$ after *post hoc* analysis. (**a**) *Controlling Arm* and *Target* interaction for length of major axis (reach direction) of the variable error ellipses. (**b**) *Controlling Arm* and *Target* interaction for length of minor axis (perpendicular to reach direction) of the ellipses

**Detailed Results on Timing**

Reaction time, movement time, total time, time to peak velocity, and time after peak velocity were assessed to identify patterns and differences in the timing of participants' reaches. Surprisingly, the results showed that reaction times were longer for the controlling right arm than the left arm (Supplementary Fig. 8a). This observation was supported by a three-factor ANOVA on the data for reaction time, which identified a main effect of *Controlling Arm* [$F(1, 35) = 4.2$, $p < 0.05$, $\eta^2_p = 0.11$]. One possibility is that participants may have taken slightly longer before initiating right-arm reaches because they were preparing a more precise movement. Another possibility is that the right arm was more sensitive to the visual identity of the virtual arm, where participants may have required more time to process the visual arm information before initiating movement. A main effect of *Target* [$F(7, 245) = 8.3$, $p < 0.001$, $\eta^2_p = 0.19$] was also identified.

In contrast, movement times were shorter for right-arm reaches than left-arm reaches (Supplementary Fig. 8b). In support of this observation, a three-factor ANOVA on the data for movement time identified a main effect of *Controlling Arm* [$F(1, 35) = 4.6$, $p < 0.05$, $\eta^2_p = 0.12$]. The ANOVA also revealed an interaction between *Controlling Arm* and *Target*, $F(7, 245) = 5.0$, $p < 0.001$, $\eta^2_p = 0.12$, and a main effect of *Target* [$F(7, 245) = 25.1$, $p < 0.001$, $\eta^2_p = 0.42$]. Summing together reaction time and movement time, total time appeared to have neither a main effect of *Controlling Arm* nor *Virtual Arm* (Supplementary Fig. 8c). A three-factor ANOVA on the data for total time identified an interaction between *Controlling Arm* and *Target* [$F(7, 245) = 2.8$, $p < 0.01$, $\eta^2_p = 0.07$] and a main effect of *Target* [$F(7, 245) = 15.4$, $p < 0.001$, $\eta^2_p = 0.31$].

Similarly, time to peak velocity showed no main effects (Supplementary Fig. 9a). A three-factor ANOVA on the data for time to peak velocity identified an interaction between *Controlling Arm* and *Target* [$F(7, 245) = 18.3$, $p < 0.001$, $\eta^2_p = 0.34$] and a main effect of *Target* [$F(7, 245) = 60.3$, $p < 0.001$, $\eta^2_p = 0.63$]. Time after peak velocity showed the same general pattern as movement time, with shorter times for the controlling right arm than the left arm (Supplementary Fig. 9b). This was supported by a three-factor ANOVA on the data for time after peak velocity, which identified a main effect of *Controlling Arm* [$F(1, 35) = 10.0$, $p < 0.01$, $\eta^2_p = 0.22$]. One possible explanation is that more uncertainty due to using the less dextrous left arm might cause slower movements at the end of a reach. Another possible explanation is that since participants showed a general pattern of overreaching more during left arm movements than right arm movements, it might have taken a bit more time to complete each reach with the left arm compared to the right. An interaction between *Controlling Arm* and *Target*, $F(7, 245) = 2.3$, $p < 0.05$, $\eta^2_p = 0.06$, and a main effect of *Target*, $F(7, 245) = 10.9$, $p < 0.001$, $\eta^2_p = 0.24$ were also identified. As shown in Supplementary Fig. 10, the effect of controlling arm across all timing variables that interacted with *Target* only varied in magnitude and not direction whereby the controlling right arm showed faster times than the controlling left arm. The exception was time to peak velocity, where each arm appeared slower for targets on its own side of space (Supplementary Fig. 10c). This may reflect direction-specific biomechanical demands rather than a consistent timing advantage for one arm.

Since timing variables only depended on the *Controlling Arm* and not the *Virtual Arm*, these results suggest that the effects of the *Virtual Arm* on accuracy variables were unlikely to be driven by a speed-accuracy trade-off.

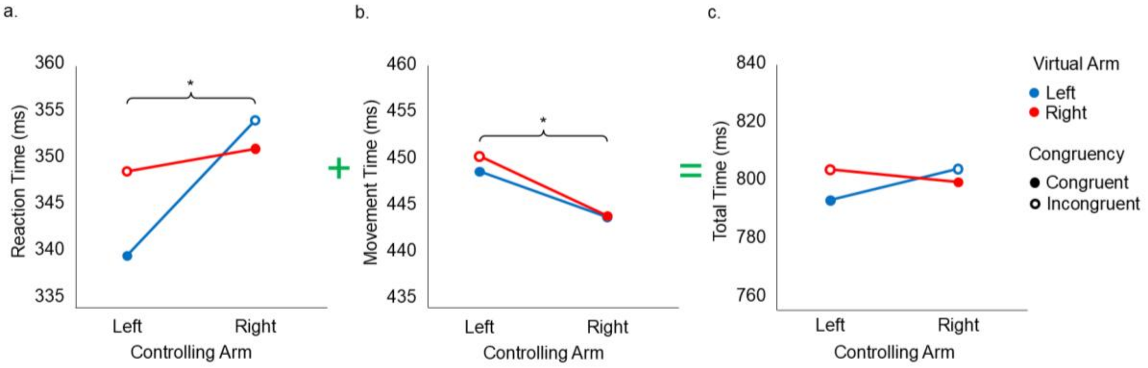


**Supplementary Fig. 8** Scores on reaction time, movement time, and total time across conditions. Main effects are shown with braces. *$p < 0.05$, **$p < 0.01$, ***$p < 0.001$. (**a**) Median reaction time. A main effect of *Controlling Arm* was identified. (**b**) Median movement time. A main effect of *Controlling Arm* was identified. (**c**) Median total time

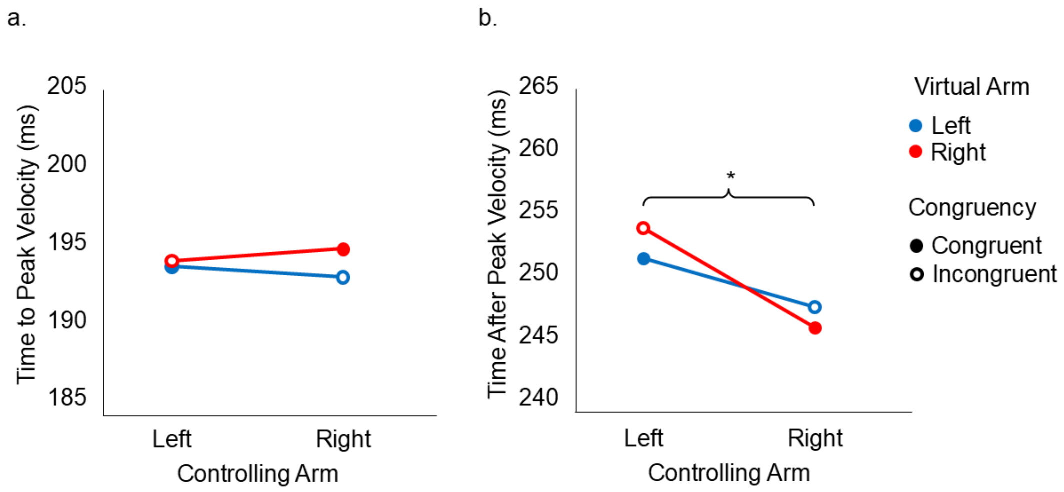


**Supplementary Fig. 9** Scores on timing dependent variables across conditions. Main effects are shown with braces. *$p$ < 0.05, **$p$ < 0.01, ***$p$ < 0.001. (**a**) Median time to peak velocity. (**b**) Median time after peak velocity. A main effect of *Controlling Arm* was identified

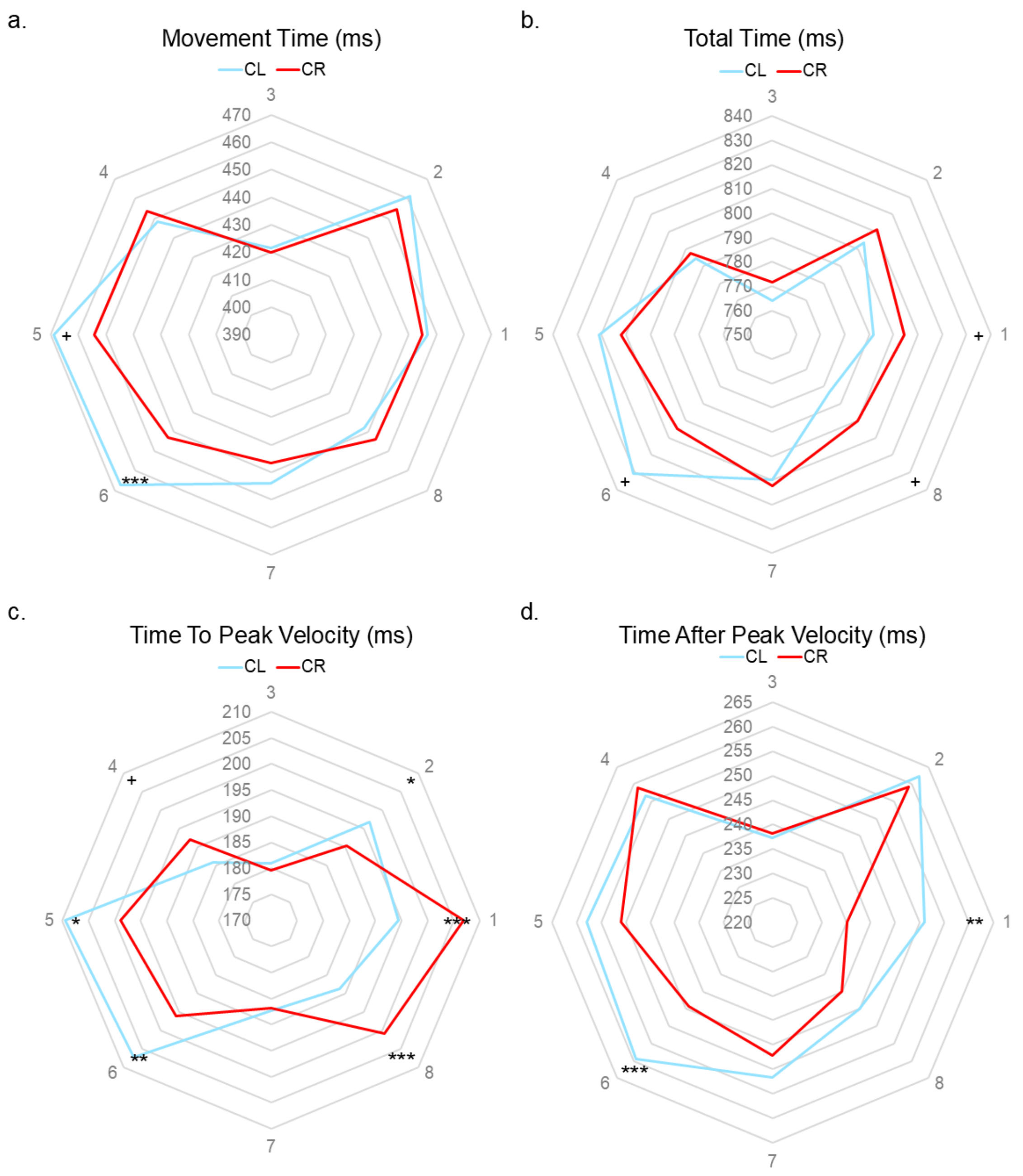


**Supplementary Fig. 10** *Controlling Arm* and *Target* interactions for timing scores. + $p_{\text{uncorr}} < 0.05$, ++ $p_{\text{uncorr}} < 0.01$. *$p < 0.05$, **$p < 0.01$, ***$p < 0.001$ after *post hoc* analysis. (**a**) *Controlling Arm* and *Target* interaction for median movement time. (**b**) *Controlling Arm* and *Target* interaction for median total time. (**c**) *Controlling Arm* and *Target* interaction for median time to peak velocity. (**d**) *Controlling Arm* and *Target* interaction for median time after peak velocity
